# The rate and spectrum of mosaic mutations during embryogenesis revealed by RNA sequencing of 49 tissues

**DOI:** 10.1101/687822

**Authors:** Francesc Muyas, Luis Zapata, Roderic Guigó, Stephan Ossowski

**Affiliations:** Institute of Medical Genetics and Applied Genomics, University of Tübingen, Tübingen, Germany; Center for Genomic Regulation, The Barcelona Institute of Science and Technology, Barcelona, Spain; Universitat Pompeu Fabra (UPF), Barcelona, Spain; Centre for Evolution and Cancer, The Institute of Cancer Research, London, UK

**Author notes:** Corresponding authors: Stephan Ossowski, Francesc Muyas.

**Keywords:** **Genetic** mosaicism, human embryogenesis, mosaic mutation rate

## Abstract

**Background:** Mosaic mutations acquired during early embryogenesis can lead to severe early-onset genetic disorders and cancer predisposition, but are often undetectable in blood samples. The rate and mutational spectrum of embryonic mosaic mutations (EMMs) have only been studied in few tissues and their contribution to genetic disorders is unknown. Therefore, we investigated how frequent mosaic mutations occur during embryogenesis across all germ layers and tissues.

**Results:** Using RNA sequencing data from the Genotype-Tissue Expression (GTEx) cohort comprising 49 normal tissues and 570 individuals, we found that new-borns on average harbour 0.5 - 1 EMMs in the exome affecting multiple organs (1.3230 × 10^−8^ per nucleotide per individual), a similar frequency as reported for germline *de novo* mutations. Our multi-tissue, multi-individual study design allowed us to distinguish mosaic mutations acquired during different stages of embryogenesis and adult life, as well as to provide insights into the rate and spectrum of mosaic mutations. We observed that EMMs are dominated by a mutational signature associated with spontaneous deamination of methylated cytosines and the number of cell divisions. After birth, cells continue to accumulate somatic mutations, which can lead to the development of cancer. Investigation of the mutational spectrum of the gastrointestinal tract revealed a mutational pattern associated with the food-borne carcinogen aflatoxin, a signature that has so far only been reported in liver cancer.

**Conclusion:** In summary, our multi-tissue, multi-individual study reveals a surprisingly high number of embryonic mosaic mutations in coding regions, implying novel hypotheses and diagnostic procedures for investigating genetic causes of disease and cancer predisposition.

## Background

Genetic mosaicism describes the co-existence of genetically different cell populations in an individual developing from a single fertilized egg (Youssoufian and Pyeritz, 2002; Biesecker and Spinner, 2013; Acuna-Hidalgo et al., 2016). Mosaicism has been associated with a broad range of genetic diseases (Campbell et al., 2015), including neurological disorders (Poduri et al., 2013; Halvorsen et al., 2016), brain malformation and overgrowth syndromes (Lindhurst et al., 2011; Rivière et al., 2012), autism spectrum disorders (Yurov et al., 2007), and cancer predisposition syndromes (Prochazkova et al., 2009; Ruark et al., 2013). Mosaicism can lead to genetic disorders that are embryonic lethal when occurring in germ cells (Happle, 1987), or result in a milder phenotype than a constitutive mutation (Plant et al., 2000). The timing of mutations during embryogenesis (e.g. cleavage, blastulation, implantation, gastrulation, neurulation and organogenesis) influences the fraction of affected cells and organs in the organism (Campbell et al., 2015; Acuna-Hidalgo et al., 2017). Moreover, when occurring during gametogenesis mosaic mutations can be passed on constitutionally to multiple offspring (Acuna-Hidalgo et al., 2016).

As expected, mosaic mutations are found in the form of single nucleotide variants (SNVs), insertions and deletions (indels) and copy number variants (CNVs), and have been studied using array technology (Pham et al., 2014) as well as Next Generation Sequencing (NGS) (Huang et al., 2014; Acuna-Hidalgo et al., 2015). A SNParray-based study of the Children’s Hospital of Philadelphia found that 17% of the diagnosed cases were caused by mosaic aneuploidies (Conlin et al., 2010). Acuna-Hidalgo and colleagues suggested that around 7% of presumed germline de novo mutations are in fact post-zygotic mosaic mutations (Acuna-Hidalgo et al., 2015). Using whole-genome sequencing of normal blood from 241 adults Ju et al. (Ju et al., 2017) estimated that approximately three mutations are accumulated per cell division during early embryogenesis. However, despite their potential importance for human disease, previous studies of mosaic mutations have focused on only one or few tissues or organs, e.g. using whole exome sequencing data of brain tissues (Wei et al., 2018a) or blood (Acuna-Hidalgo et al., 2015). Therefore, a comprehensive view of mosaic mutations arising during embryogenesis, including their rate and mutational spectrum, is missing. Here, we exploit 10,097 RNA-seq samples from 49 different tissues and 570 individuals of the Genotype-Tissue Expression (GTEx) cohort (Lonsdale et al., 2013) to uncover the rate and spectrum of mosaic mutations acquired post-zygotic during early embryogenesis.

## Results

### Somatic variant calling in RNA-seq data

Somatic variant detection using RNA-seq data is challenging, especially if subclonal mutations with allele fractions as low as 5% are of interest (Yizhak et al., 2019). We therefore developed a highly accurate multi-sample variant calling procedure, which models nucleotide-specific errors, removes germline variants and confounders such as RNA editing sites, and generates a multi-individual, multi-tissue call matrix (3D genotype matrix) by re-genotyping potentially variable sites across thousands of GTEx RNA-seq samples (Figure 1a). We trained a random forest classifier (*RF-RNAmut*) to distinguish true from false variant calls using whole exome sequencing (WES) and RNA-seq data from the ICGC Chronic Lymphocytic Leukaemia project (Puente et al., 2015) for generating training and test data (Methods). High confidence somatic variant calls with >0.15 VAF in tumour WES data were identified in RNA-seq with 71% sensitivity and 85% precision, and sensitivity was positively correlated with VAF (Supp. Table 1). Germline variants found in tumour and normal WES data, although not our interest in this study, were identified in RNA-seq data with 86% sensitivity and 95% precision.

**Figure 1.**
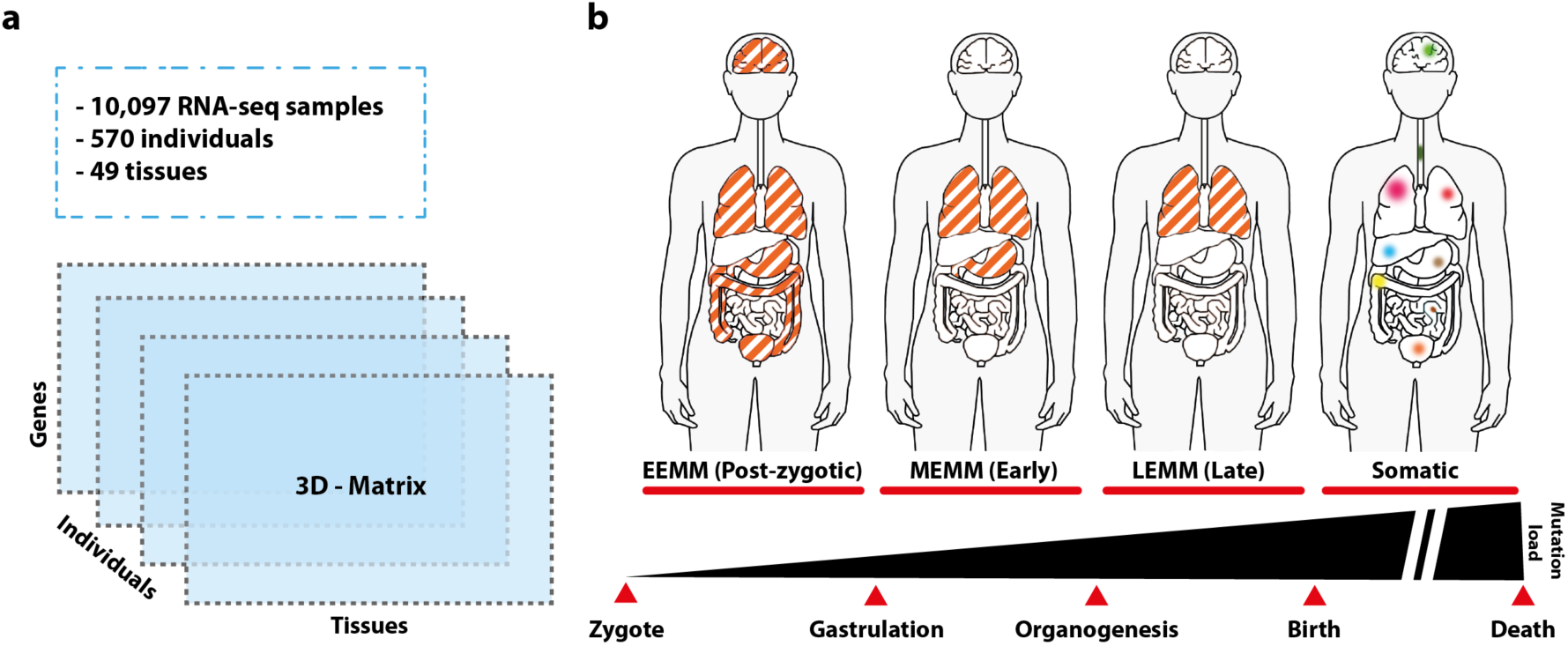
Identification of mosaic mutations acquired during various developmental stages and adult life. a) 10,097 RNA-seq samples from 49 tissues and 570 individuals (GTEx release 7) were used to generate a 3D genotype matrix, which facilitated the identification mosaic mutations and determination of their germ layer or organ of origin. b) Definition of mosaic mutation types depending on the developmental stage during which they occur: early-embryonic mosaic mutations (EEMMs) occurring during the first few cell divisions of the zygote until implantation of the embryo, mid-embryonic mosaic mutations (MEMMs) acquired during gastrulation or neurulation (example in image: mutation in endoderm), late embryonic mosaic mutations (LEMMs) acquired during early organogenesis, and somatic mutations acquired after birth. See also Supp. Figure 1 for the embryogenesis lineage tree used in the study.

### Rate and spectrum of early mosaic mutations during embryogenesis

In order to identify mosaic mutations acquired during embryonic development, we computed the 3D genotype matrix for 9,704 samples of the GTEx cohort comprising 526 cancer-free individuals and 49 tissues (see Methods for sample selection criteria). We contrasted the 3D genotype matrix with the embryogenesis lineage tree (Supp. Figure 1) to identify the most likely germ layer or tissue of origin of each mutation. We first removed variants occurring in all expressed tissues with average VAF greater 0.35, as they might constitute *de novo* germline variants. Then, we defined three types of embryonic mosaic mutations (EMMs): early (pre-implantation), mid (gastrulation and neurulation) and late (organogenesis) embryonic mosaic mutations (EEMMs, MEMMs, LEMMs in Figure 1b). EEMMs appeared during the first few divisions of the zygote (cleavage, blastulation, implantation) and therefore are present in the Ectoderm and Mesendoderm (Mesoderm and/or Endoderm). MEMMs are mutations found in at least two tissues of the same individual that originate from the same germ layer. We define LEMMs as mutations present in a large cell fraction of a single organ, which are not the consequence of somatic clonal expansions. Finally, we also screened for postnatal and adult somatic mutations in the transcriptome of all cancer-free individuals.

To minimize false negatives we focused our analysis on housekeeping genes constitutively expressed in the majority of tissues and samples (7,630 genes with TPM > 5 in at least 75% of tissues). After strict filtering (Methods), we identified 58 putative EEMMs and 37 MEMMS in 7630 constitutively expressed genes. We estimated a rate of 8.1164 × 10^−9^ (CI (95%) = [7.0973 × 10^−9^ to 9.1292 × 10^−9^]) EEMMs and a rate of 5.1166 × 10^−9^ (CI (95%) = [4.5592 × 10^−9^ to 5.6740 × 10^−9^]) MEMMs per nucleotide and individual for exonic regions. Following an approach for extrapolating tumour mutation burden (TMB) from gene panels to exomes (45Mbp exonic regions) (Chalmers et al., 2017), we estimated a mean of 0.37 exonic EEMMs (Figure 2a) and 0.23 exonic MEMMs (Figure 2b) per individual (0.44 and 0.275 when correcting for precision and sensitivity of our variant calling algorithm). Using different thresholds for constitutively expressed genes only marginally affected the estimated rate of EEMMs or MEMMs (Figure 2a-b, Supp. Table 2).

**Figure 2.**
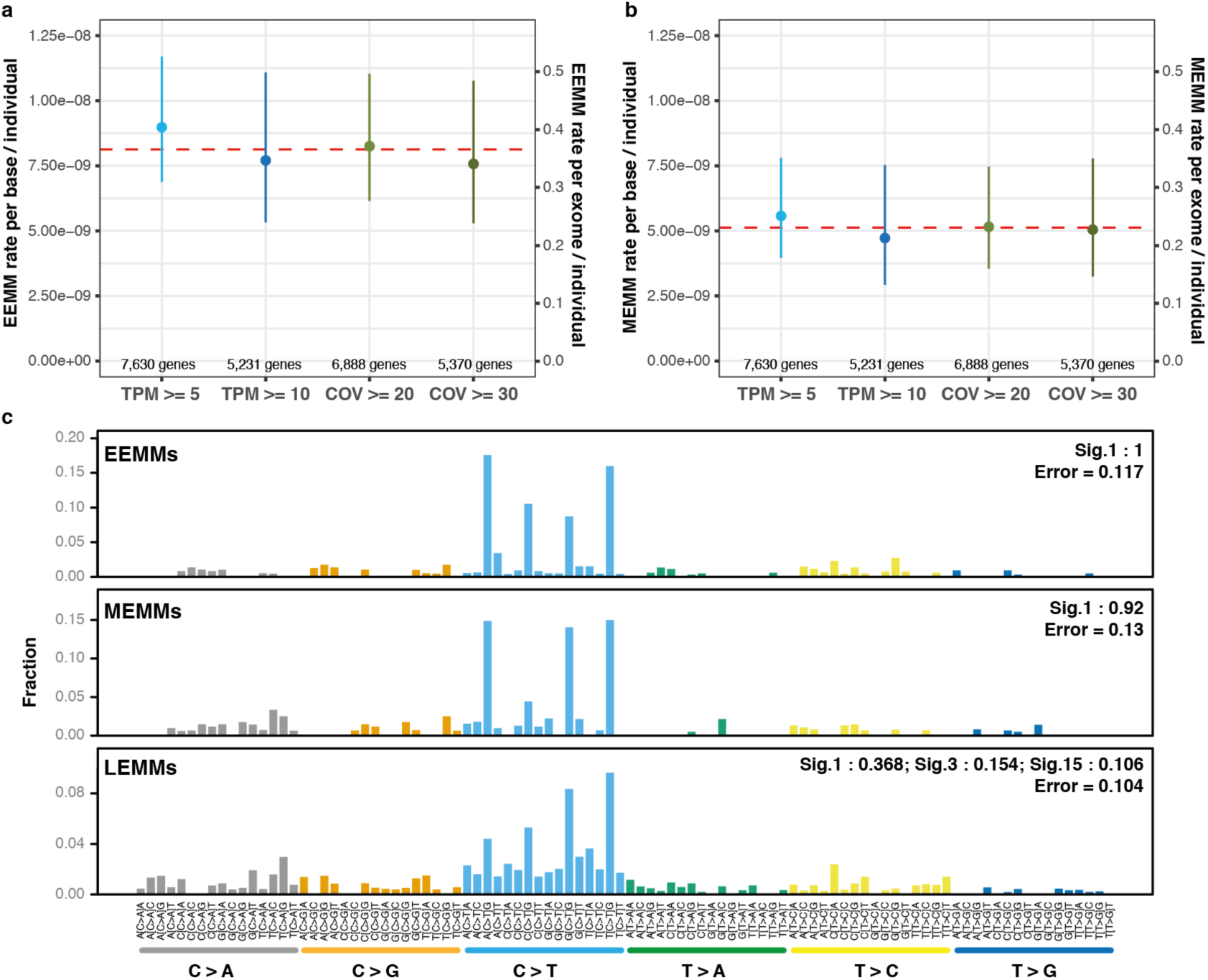
Rate and mutational signatures of mosaic mutations in healthy individuals acquired during embryogenesis. For a) and b), the left Y-axis represents the mutational rate per nucleotide, the right Y-axis represents the extrapolated number of mosaic mutations expected in 45 Mbps coding exons, and the dashed red line indicates the mean rate/number between different parameter setting (i.e. different definitions of constitutively expressed genes). a) Rate of early-embryonic mosaic mutations (EEMMs) acquired during the first few divisions of the zygote. We estimated a mean rate of EEMMs per base and individual of 8.1164 × 10^−9^ (CI (95%) = [7.0973 × 10^−9^ to 9.1292 × 10^−9^]). b) Mid-embryonic mosaic mutations (MEMM) affecting at least 2 tissues. We estimated a mean of 5.1166 × 10^−9^ MEMMs per nucleotide and individual (CI (95%) = [4.5592 × 10^−9^ to 5.6740 × 10^−9^]). c) Mutational signature of early-(EEMMs), mid-(MEMMs) and late-embryonic mosaic mutations (LEMMs).

On average, a specific EEMM was detectable in 63.6% of the tissues of an individual expressing the respective gene, consistent with the assumption that they arose during the first divisions of the zygote. Interestingly, only 41% of EEMMs in genes expressed in blood were detectable in blood samples, which could be explained by the asymmetric cell doubling model (unequal contribution of early embryonic cells to adult somatic tissues) suggested by Jue *et al* (Ju et al., 2017). Hence, a large fraction of mosaic mutations would be missed by blood-based genetic diagnostic tests. As expected, we observed a positive correlation between the variant allele fraction of EEMMs/MEMMs and the number of tissues supporting the variant (Rho=0.56; p-value=3.24 × 10^−9^). Moreover, mutations occurring earlier in development also showed a greater proportion of cells carrying the variant (Rho=-0.39; p-value=7.83 × 10^−5^, Supp. Figure 2).

The combined rate of EEMMs and MEMMs of 1.32 x× 10^−8^ is comparable to the estimated rate of *de novo* germline mutations reported in the literature (Acuna-Hidalgo et al., 2015, 2016), ranging from 1.0 to 1.8×10^−8^ per nucleotide per generation (44 to 82 mutations per genome (Acuna-Hidalgo et al., 2016), or ∼0.5 – 1 mutations per exome (45Mbp) per individual). Recently, several genetic disease studies indicated that more than 50% of sporadic cases can be explained by *de novo* germline mutations (Poduri et al., 2013; Acuna-Hidalgo et al., 2016). Consequently, embryonic mosaic mutations are similarly likely to explain a significant fraction of sporadic genetic disease cases, and a substantial fraction of germline *de novo* variants identified in blood are potentially postzygotic mutations. Moreover, we likely underestimated the rate of EEMMs and MEMMs due to factors such as allele specific expression, nonsense-mediated decay, and more effective transcription-coupled repair in highly expressed genes. As most of the disease-causing mosaic mutations cannot be detected by sequencing blood-derived DNA, these variants have likely been missed in past studies, and could explain a substantial part of the missing heritability.

In order to identify the most likely processes causing early- and mid-embryonic mosaic mutations we investigated their mutational signatures. We found that a large fraction of EEMMs and MEMMs (1 and 0.92) could be explained by Signature 1 (Alexandrov et al., 2013, 2015; Nik-Zainal et al., 2016) (Figure 2c). Signature 1 is thought to be the result of an endogenous mutational process initiated by spontaneous deamination of 5-methylcytosine leading to C>T transitions at CpG dinucleotides and likely reflects a cell-cycle-dependent mutational clock (Alexandrov et al., 2015). Hence, our findings indicate that most early- and mid-embryonic mosaic mutations occur spontaneously with very limited contributions from exposure to environmental factors or other endogenous processes. Furthermore, our results clearly distinguish early mosaic mutations from germline *de novo* mutations, which are dominated by Signature 5 characterised by A>G transitions (Acuna-Hidalgo et al., 2016).

### Late embryonic mosaic mutations arising during organogenesis

Our definitions of EEMMs and MEMMs prevent identification of organ-specific mutations acquired during organogenesis. We therefore screened for late embryonic mosaic mutations (LEMMs, Figure 1b), which we defined as tissue-specific mutations at high cell fraction (VAF >= 0.2). Here, we excluded tissues previously shown to be affected by clonal expansion of mutated cells such as esophagus-mucosa, sun-exposed skin (Martincorena et al., 2015, 2018; Chalmers et al., 2017; Yizhak et al., 2019; Yokoyama et al., 2019) and whole blood (Acuna-Hidalgo et al., 2017), which also showed the highest somatic mutation rates in our analysis (Supp. Figure 3). We identified 377 mutations across all individuals, considering any gene expressed in at least one tissue, resulting in an estimate of 2.44 × 10^−9^ (CI [0.95] = [1.86 × 10^−9^ - 3.03 × 10^−9^]) LEMMs per nucleotide per tissue per individual, and extrapolating to 0.11 (CI [0.95] = [0.084 - 0.137]) mutations per exome per tissue. Notably, the average rate of LEMMs (2.23 × 10^−9^) for brain tissues closely resembled the estimates by Wei *et al.* (Wei et al., 2018b) (2.55 × 10^−9^) obtained using WES data of brain tissues. In sum across all 43 examined tissues we estimated 4.7 LEMMs per exome per individual. However, LEMMs are indistinguishable from mutations in clonal expansions acquired after birth (Martincorena et al., 2015, 2018; Yizhak et al., 2019), and the rate of LEMMs is therefore likely overestimated. Nonetheless, our results indicate that organ-specific mosaic mutations arising during organogenesis could significantly contribute to the phenomenon of missing heritability in rare genetic diseases as well as cancer predisposition.

**Figure 3.**
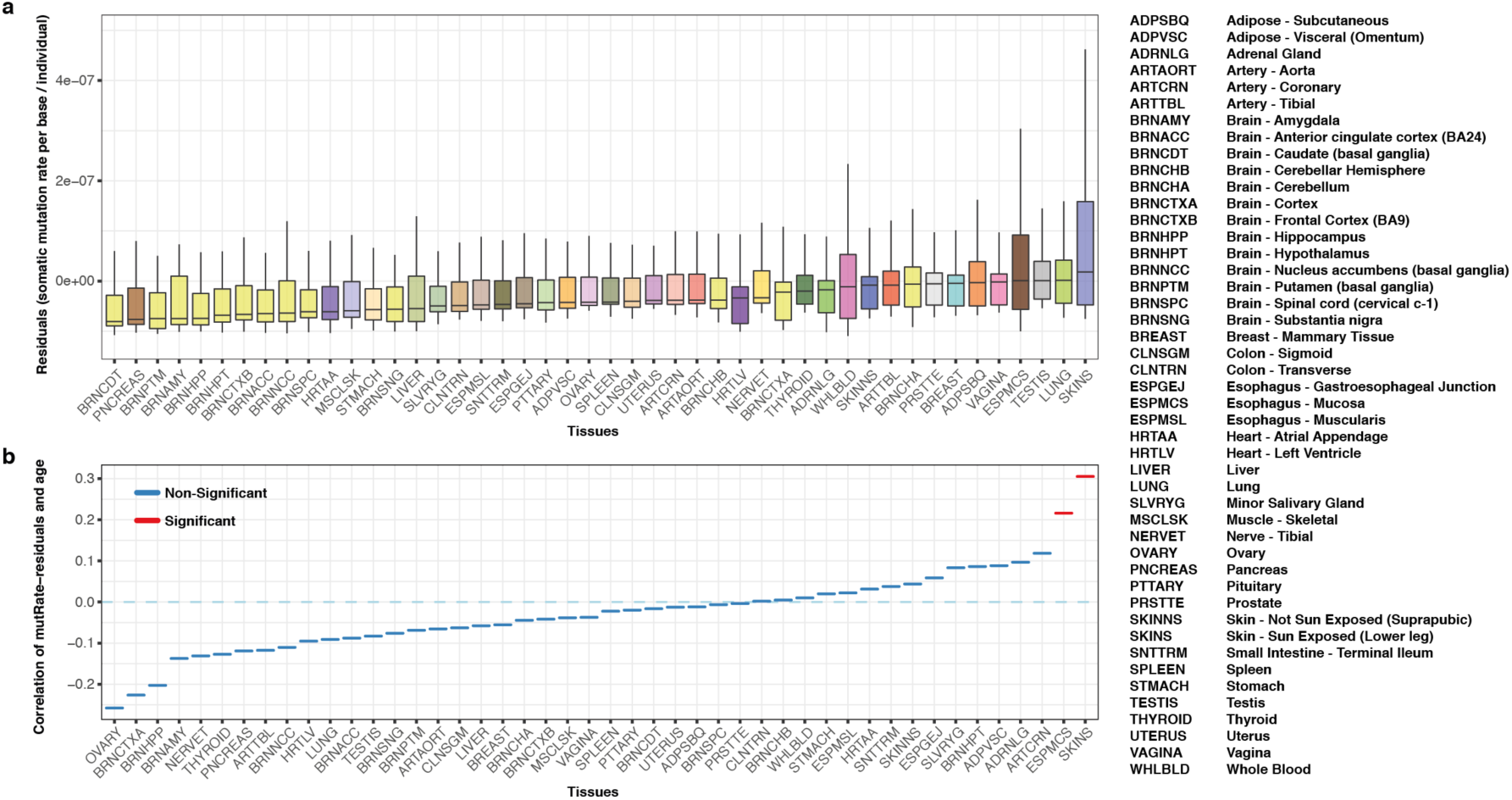
Rate of somatic mutations varies significantly across the 46 tissues of the GTEx cohort. (ignoring Kidney, Cells-EBV-transformed lymphocytes and Cells-transformed fibroblasts for technical reasons, see Methods). a) Distribution of residuals of the somatic mutation rate per base and individual residuals (mutRate-residuals) across analysed tissues. MutRate-residuals represent the somatic mutation rates corrected for non-biological confounders such as PCR duplication rate, RIN, cohort, and read coverage. b) Spearman correlation between mutRate-residuals and age per tissue. Colours show the significance of the correlation test after FDR correction (q-value < 0.05 in red).

### Rate and mutational signatures of tissue-specific somatic mutations

To identify other mutation processes leading to the accumulation of somatic mutations during adult life, we next studied mutational signatures across all tissue-specific somatic variants identified in the GTEx cohort. Considering only variants with VAF >= 0.05, we identified 8,780 somatic mutations in 8,351 samples representing 46 tissues (Methods and Supp. Figure 4). After removal of technical confounders (PCR duplicate rates, RIN, TRISCHD, coverage, laboratory), we observed the highest mutation burden for sun-exposed skin, lung, testis, esophagus-mucosa and vagina (Figure 3a and Supp. Figure 5), as previously reported (Yizhak et al., 2019). As expected, sun-exposed skin showed significantly higher mutation burden than non-sun-exposed skin, while brain tissues showed the lowest somatic mutation burden. Finally, we tested if residual mutation rates were related with age of individuals for each tissue individually (Figure 3b). Only two tissues showed a significant association between age and mutational rates (after FDR correction), namely sun-exposed skin (Rho = 0.31; qval = 1.19 × 10^−7^) and esophagus-mucosa (Rho = 0.22; qval = 2.82 × 10^−3^), as previously reported (Martincorena et al., 2015, 2018; Yokoyama et al., 2019). Using dN/dS as a measure of selection, we observed a lack of selection in highly-expressed genes at a pan-tissue level (dN/dS = 0.98, CI[95] = [0.92 – 1.06]). However, when focusing on cancer genes we observed strong positive selection for sun-exposed skin and esophagus-mucosa (Supp. Figure 6). Mutations in *NOTCH1* and *TP53* disproportionally contributed to the high dN/dS values and showed the highest overall mutation rates. *NOTCH1* showed stronger positive selection than *TP53* in both esophagus-mucosa and skin sun-exposed (dN/dS of 8.46 vs. 4.57 and dN/dS of 4.01 vs. 2.85, respectively, Supp. Table 3). Interestingly, we did not find positive selection of these two genes in any other tissues, and no other gene reached significance in any of the tissues.

**Figure 4.**
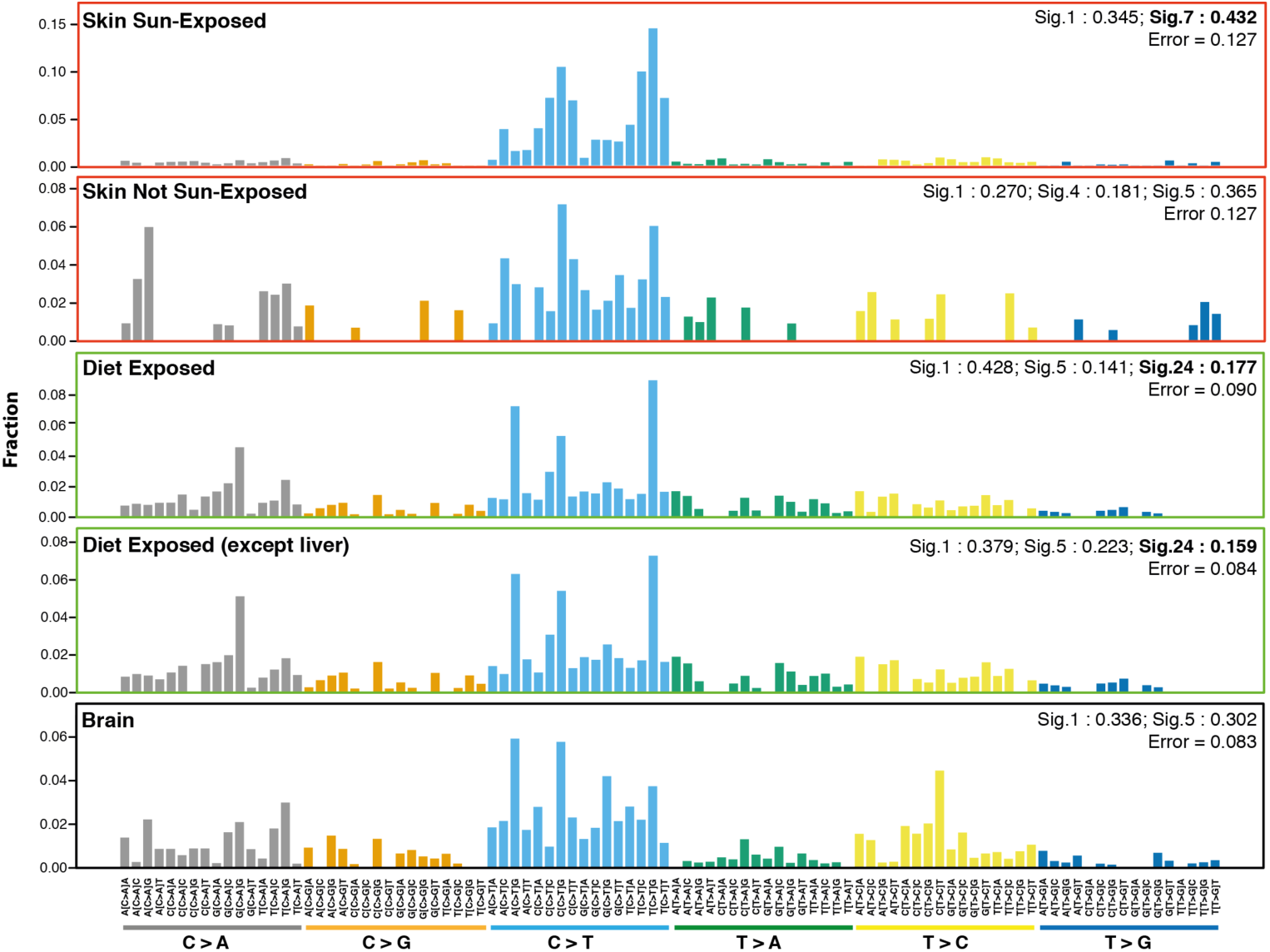
Mutational signatures observed in Skin sun-exposed, Skin not sun-exposed, Diet exposed tissues, Diet exposed (except liver) tissues, and Brain tissues. (considered control tissues with low exposure to environmental factors). Detailed descriptions of signatures are available at https://cancer.sanger.ac.uk/cosmic/signatures_v2.

### Aflatoxin mutational signature in organs of the dietary tract

Previous studies have analyzed the spectrum of somatic mutations in healthy esophagus and skin (Martincorena et al., 2015, 2018; Yizhak et al., 2019; Yokoyama et al., 2019), identifying mutational signatures (Alexandrov et al., 2013) 1, 5 and 7 (https://cancer.sanger.ac.uk/cosmic/signatures_v2). Our analysis of mutational signatures for patients who died at advanced age (>=60 years old) revealed that ultraviolet-light (UV) exposure (signature 7) was predominant in sun-exposed skin, while it was absent from non-sun-exposed skin (Figure 4). Interestingly, studies of the mutational signatures found in healthy tissues forming the dietary tract are lacking, although the constant exposure to food likely leads to a particular mutational spectrum. We therefore performed a pan-gastrointestinal-tract mutational signature analysis considering colon, esophagus-mucosa, liver, small intestine and stomach. Apart from signatures 1 and 5, which are frequently observed in most tissues, we found a signature explained by the mutagenic effect of dietary aflatoxin (Signature 24). The aflatoxin signature explained a fraction of 0.18 of the mutational spectrum in the tissues of the gastrointestinal tract (Figure 4). Furthermore, we saw a strong enrichment of the characteristic CGN > CTN mutations not observed in any other tissue.

Aflatoxin B_1_ (AFB1) is a potent mutagen and carcinogen typically found in grains contaminated with the food spoilage fungus, *Aspergillus flavus*. Dietary exposure to aflatoxin B_1_ (AFB1) is a known risk factor for human hepatocellular carcinoma (HCC), the third leading cause of cancer death worldwide. One of aflatoxins degradation products, the metabolite exo-epoxide, forms a covalent bond with guanyl N7 (AFB1-N7-Gua), ultimately leading to G>T mutations during replication. Consistently, signature 24 has previously been found in a subset of liver cancers (Chawanthayatham et al., 2017; Zhang et al., 2017), but has not been reported for other cancer entities. We therefore tested, if the observed enrichment of signature 24 was solely introduced by a strong mutagenic effect in the liver. To the contrary, when excluding liver from the analysis, the aflatoxin signature was still found at a similar level, explaining a fraction of close to 0.16 of the mutational spectrum. In comparison, we did not identify signature 24 in any other tissue (Figure 4), These results indicate that aflatoxin-related mutations are frequent in all tissues of the gastrointestinal tract, and might play a role in the development of cancer in several organs.

## Discussion

The accumulation of DNA mutations during life is inevitable, despite the many cellular mechanisms involved in the preservation of genome integrity. In this study, we presented a novel analysis strategy using RNA-seq data of multiple tissues per individual to identify mosaic mutations occurring during various stages of embryo development. Using the human embryogenic lineage tree, we approximated the time point of the mutation events as well as the affected germ layer or developing organ. We demonstrated how to distinguish, to some extent, embryonic mosaic mutations from de novo germline mutations and somatic mutations in clonal expansions acquired after birth.

Analysing RNA sequence data from 49 tissues and 570 patients we found that new-borns on average harbour 0.5 - 1 mosaic mutation in coding exons affecting multiple tissues and organs, and likely an even larger number of organ-specific coding mutations. Postzygotic and early embryonic mosaic mutation patterns are dominated by signature 1, which is associated with aging and cell division. Hence, they largely result from spontaneous deamination of methylated cytosines without showing any influence of external mutagens. Moreover, our estimates suggest that embryonic mosaic mutations are as frequent as germline *de novo* mutations and could explain a substantial fraction of unresolved cases of sporadic and rare genetic diseases, as well as play a role in cancer predisposition.

The recognition of a widespread and under-recognised role of mosaic mutations in genetic disease would have many implications for genetic diagnostics procedures (Lupski, 2013). We have furthermore demonstrated that a substantial fraction of EMMs is not detectable in blood cells, a finding which has important implications for clinical diagnostics, as samples from the affected tissue are often unavailable. Therefore, circulating cell-free DNA could be an unbiased source for detecting mosaic mutations in any tissue.

Interestingly, our method also revealed a strong signature of the food poison aflatoxin detectable in all organs of the dietary tract. Aflatoxin mutations have previously been associated to liver cancer. Our results indicate that the role of aflatoxins in cancer development might be more widespread than previously appreciated, affecting the mutation spectrum of tumours in colon, esophagus-mucosa, liver, small intestine and stomach.

## Conclusion

In this study, based on a multi-tissue, multi-individual analysis, we found a surprisingly high number of embryonic mosaic mutations in exonic regions of healthy individuals, implying novel hypotheses and diagnostic procedures for investigating genetic causes of disease, cancer predisposition and ageing.

## Methods

### Samples

In this study we used release 7 of the Genotype-Tissue Expression (GTEx) (Lonsdale et al., 2013) project (dbGaP accession phs000424.v7.p2), including RNA-seq data for 49 tissues from 570 individuals. We included only individuals for which Whole Genome Sequencing (WGS) data was available (necessary for distinguishing somatic from germline variants) and for which at least 8 tissues were analyzed by RNA-seq. Furthermore, we only included tissues for which RNA-seq data from at least 25 donors was available. Filtering by these criteria resulted in RNA-seq data from 10,097 samples distributed over 570 individuals and 49 tissues. Additional QC and filtering steps were performed depending on the specific analysis, as detailed below.

### Pipeline for somatic variant prediction in RNA-seq data

Reads were aligned using STAR (version 2.7) against the human reference genome (GRCh37) and the resulting BAM files were post-processed in order to remove alignment artefacts. PCR duplicates were marked using Picard (version 2.10.1) and reads mapping to different exons were split using SplitNCigar (part of GATK 3.7 package). Additionally, reads not overlapping with annotated human exons (ENSEMBL GRCh37 release 95) or aligning to immunoglobulin genes (potentially hyper-mutated) were removed from downstream analysis. Furthermore, reads aligning with mapping quality lower than 255, more than one gap opening or more than 3 mismatches were filtered. Finally, in order to avoid systematic alignment errors at the extremes of the reads (which also include the ‘inner ends’ of reads split across introns, i.e. breakpoints of spliced-reads), we trimmed the first and last 4 bases from each read-end or read-breakpoint (BamUtil version 1.0.14).

Using the post-processed BAM files, we computed a three-dimensional genotype-matrix (variant x tissue x individual) for all positions found to have a significant alternative allele call in at least one sample. This algorithm consists of two main steps:

Step 1: Single sample variant calling. First, base counts are obtained with *samtools mpileup* (version 1.3.1) followed by post-processing using custom scripts (Python and R scripts). We modeled the error rate (ER) distribution for each sample using a beta-binomial distribution. Counts of alternative (non-reference) reads at homozygous-reference positions (germline) are distributed following a binomial distribution with parameter P (error rate), which is a random variable that follows a Beta distribution with parameters *α* and *β*.

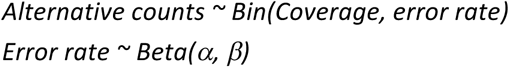

As the error rate differs depending on the nucleotide change (for example due to DNA oxidation artifacts affecting only a specific base), we modeled error distributions independently for each possible nucleotide change (A>C, A>T, A>G, C>A, C>T, C>G). Finally, we identified all sites showing alternative allele counts significantly deviating from the ER distribution after FDR correction. Additional filtering criteria were applied for each site, including a minimum alternative allele count of 4 (each having at least base quality of 20), minimum read coverage of 10, alternative calls presented in forward and reverse strand following the same distribution as for reference counts (i.e. no strand bias), variant allele frequency (VAF) greater or equal to 5%, and minimum distance of 20 bp between variable sites in the same sample.

Step 2: Multi-sample re-calling of all potentially variable sites across all individuals and tissues is performed using a custom algorithm in order to build the three-dimensional genotype matrix. To this end, sites passing step 1 as significant in at least one sample were evaluated in each sample using the beta-binomial distribution as described for single samples, but with less stringent post-filtering criteria, resulting in one of four possible filter states per sample: NO_EXPRESSION, HOM_REF, LOW_QUALITY or PASS. Furthermore, the exact reference-like and alternative allele counts are stored in the Coordinates x Tissue x Individual matrix.

### A random forest model for multi-tissue, multi-individual germline and somatic variant calling from RNA-seq data

We next aimed at training a random forest classifier distinguishing true from false positive variant calls in RNA-seq data. To this end we selected 40 cases studied as part of the ICGC Chronic Lymphocytic Leukemia project, for which whole exome sequencing (WES) data for tumor and normal sample and RNA-seq data for tumor samples are available. RNA-seq based variant calling was performed as described above for GTEx samples. Additionally, we obtained the reference and alternative allele counts from tumor and normal WES data for all putative calls identified in RNA-seq data. Finally, we used the WES data to predicted high quality germline and somatic variant calls using GATK HaplotypeCaller and MuTect2 as described before (Zapata et al., 2017; Muyas et al., 2019).

Next, variants identified in RNA-seq data were randomly split into training and test sets for RF model training and testing, with the restrictions that:

- Training and test set contain a similar number of true and false events according to WES data
- Training and test sets have a uniform distribution of variant allele frequencies, except for variants with VAF < 10%, which were doubled (in order to increase sensitivity of the RF for low VAF)

In addition, a set of non-overlapping high quality calls from WES data was incorporated in training and test sets. We labeled as true variants any site with VAF >= 5% and at least 2 reads supporting the alternative allele in WES data, and all other sites as false variants. This procedure resulted in training and test datasets of 2402 sites each.

To train the RF model (R *randomForest* package) for distinguishing true and false positive variants (germline or somatic) called in RNA-seq data we included as features: a) alternative allele count, b) coverage, c) VAF, d) strand bias, e) blacklisted genes (Fuentes Fajardo et al., 2012), and f) average alternative base quality. As this model, termed *RF-RNAmut* from here on, returned a response value between 0 and 1 for detecting calls, we chose our cutoff based on the maximum F1 score in the training set (cutoff = 0.19). Sites with response values exceeding 0.19 were labelled as high confident variants. To finally generate the somatic mutation call set and to remove systematic calling errors we filtered variants if: (1) they were recurrently called in RNA-seq data of multiple individuals, (2) their population allele frequency in GnomAD or 1000GP was greater than 1%, (3) they overlapped with repetitive elements annotated by Repeat Masker, (4) they overlapped with low complexity regions, (5) were flagged as likely systematic analysis error by ABB (Muyas et al., 2019), or (6) they overlapped with a known RNA editing site (Kiran et al., 2012; Picardi et al., 2017; Tan et al., 2017).

We measured the performance (precision and recall) of *RF-RNAmut* + Filter on identifying a) germline and b) somatic variant calls using the test set, following the same procedure as described above. To calculate precision, we considered as true or false positive calls those variants, which were found in RNA-seq data and matched or not matched with tumor WES data, respectively. For calculating the false negative rate, we considered high-quality calls identified by MuTect2 in tumor-normal paired WES analysis that were not found in RNA-seq data. For benchmarking purposes, we only analyzed regions overlapping between RNA-seq (with more than 10x read coverage in annotated exons) and the WES enrichment kit (Agilent SureSelect 71Mb). Again, non-exonic regions, known editing sites and Immunoglobulin genes were ignored.

### Identification of mosaic mutations in the GTEx cohort

In order to obtain true mosaic variant calls for the GTEx cohort we first removed all germline variants detected by WGS analysis in any individual (GATK HaplotypeCaller) from the 3D genotype matrix. Additionally, we removed any site for which the minor allele frequency in the population was greater or equal than 1% in GNOMAD or 1000GP. Furthermore, we removed all variants present in expressed tissues of all individuals, as they likely represent systematic errors, RNA editing sites or germline de novo mutations. To further deplete calls produced by RNA-editing events (mainly A > I, less frequently C > U) we ignored known editing sites described in the literature (http://lilab.stanford.edu/ (Tan et al., 2017), found in the Darned database (https://darned.ucc.ie/download/ (Kiran et al., 2012)) or identified by the GTEx consortium (http://srv00.recas.ba.infn.it/atlas/pubs.html - REDIportal (Picardi et al., 2017)).

Next, we removed sites, which recurrently exhibit low quality (LQ) calls across multiple individuals, which are likely systematic sequencing or alignment errors. Moreover, we filtered out positions labeled as systematic errors by ABB (Muyas et al., 2019). Additionally, we removed any variant overlapping with low complexity regions or repeat regions annotated by Repeat Masker. Finally, as we did not expect mosaic mutations to be highly recurrent in different individuals, we removed sites called in more than 2 individuals of our cohort.

### Identification of early- (EEMMs) and mid -embryonic mosaic mutations (MEMMs)

In order to identify mosaic mutations acquired during early embryogenesis (cleavage, blastulation, gastrulation, neurolation and early organogenesis) we contrasted the somatic calls in the 3D genotype matrix with a lineage tree of human embryogenesis and tissue development including the 49 tissues studied here (Yu et al., 2010). In this part of the analysis, only individuals with 10 or more tissues sequenced with at least two germ layers represented by 2 sequenced tissues were included in the analysis (526 individuals). This procedure allowed us to identify mosaic mutations affecting at least two tissues, whose origin could be unambiguously mapped to a specific stage of development and/or primary germ layer.

Mosaic mutations identified in both the Ectoderm and Mesendoderm branches having zygote as most likely ancestral node, i.e. variants likely originating from the first few divisions of the zygote (cleavage, blastulation, implantation stages), were defined as early-embryonic mosaic mutations (EEMMs). In order to avoid detection of de novo germline variants as EEMMs we only considered variants with VAF less than 0.35 that were not found in all expressed tissues of an individual.

The remaining mutations found in at least two tissues of an individual were defined as mid-embryonic mosaic mutations (MEMMs) if: (1) their most likely ancestral node was not zygote; (2) they were only observed in either the ectoderm or the mesendoderm sub-tree; and (3) their appearance in the lineage tree was coherent. Contradictory (non-coherent) mutation patterns were defined as alternative alleles, which were observed in far-apart nodes in the tree, but which were undetectable in any node close to the affected tissues. In other words, mosaic mutations that required the assumption that they had occurred multiple times independently in different cells of the same individual were not considered coherent and were removed.

Finally, we defined late embryonic mosaic mutations (LEMMs) as those mutations that are restricted to one tissue/organ, but likely occurred early during organogenesis. To this end, we considered variants found in a single tissue per individual, supported by 5 or more reads and with VAF of >= 0.2. This procedure cannot distinguish mosaic mutations acquired during late embryogenesis (organogenesis) from mutations in clonal expansions acquired after birth. We therefore excluded somatic variants from tissues known to have detectable clonal expansions such as sun-exposed skin, esophagus-mucosa and whole blood.

### Estimating the rate of mosaic mutations during embryogenesis

Reliable detection of mosaic mutations in a gene using RNA-seq data and definition of the mutation’s origin in the lineage tree requires high gene expression in a majority of tissues of an individual. In order to estimate the rate of mosaic mutations we therefore focused on genes that are highly and constitutively expressed in most of the analyzed tissues. Given a large enough pool of constitutively expressed genes we can subsequently extrapolate mutation rates to the whole exome or genome, as suggested previously for measuring genome-wide tumor mutation burden (TMB) using small cancer gene panels (Chalmers et al., 2017). We used four different thresholds to define sets of constitutively expressed genes. For each set we independently estimated the rate of mosaic mutations, to ultimately evaluate the robustness of our approach by comparing the four estimates. The following definitions were used to define constitutively expressed genes:

1. Genes with TPM >= 5 in more than 75 % of all total samples (7,630 genes).
2. Genes with TPM >= 10 in more than 75 % of the total samples (5,231 genes).
3. Genes with COV >= 20 in than 75 % of the total samples (6,888 genes).
4. Genes with COV >= 30 in more than 75 % of the total samples (5,370 genes).

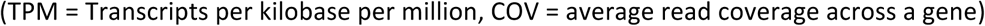

Next, we obtained all mosaic variants identified in a given set of constitutively expressed genes and calculates the number of mutations per base and individual relative to the total length of the interrogated region. Finally, we extrapolated this value to the approximate total length of all coding exons (45Mbp) in order to calculate the number of mosaic coding mutations expected on average for a newborn child. The procedure was independently performed for EEMMs and MEMMs.

For LEMMs, which were defined as tissue-specific, we considered any gene highly expressed in a given tissue of an individual (i.e. a sample). We normalized the number of mutations per base and individual relative to the interrogated region for a given sample and extrapolated this value to the approximate total length of all coding exons (45Mbp). Due to their similarity with mutations in clonal expansions the rates of LEMMs per exome per individual are likely overestimated.

### Tissue-specific somatic mutation rates

In order to study somatic mutations acquired after birth, the rate of somatic mutations, signatures of selection, as well as mutation spectra in a tissue specific manor, we performed somatic variant calling using *RF-RNAmut* without the restrictions applied for the detection of embryonic mosaic mutations. Here, we only considered somatic mutations identified in exactly one tissue per individual in order to minimize the number of mosaic mutations acquired before birth in this set. First, we performed samples-wise quality control (Supp. Figure 4) and excluded samples with the following characteristics:

- PCR duplicate rates in the top 5 %.
- Outliers for the number of callable sites (top and bottom 1% per tissue). We considered a site as callable if the read coverage was >= 10.
- Outliers for RIN (bottom 1 % per tissue)
- Outliers for mutation rate (top 1 % per tissue)
- Samples obtained from cell culture (Cells-EBV-transformed_lymphocytes, Cells-Transformed_fibroblasts)
- Individuals affected by cancer

In order to improve the statistical power, we removed tissues with less than 50 high quality samples from downstream analysis (affecting only kidney with 38 high quality samples, see Supp. Figure 4d) resulting in 8,351 samples from 46 tissues and 558 individuals. We calculated the somatic mutation rate based on the number of identified somatic mutations divided by the callable sites per sample. As quality control revealed a strong influence of technical confounders (PCR duplicate rate, RIN, average coverage, sequencing center) on the number of detectable mutations we used a linear regression model to estimate and subtract technical biases. The linear regression model uses the following variables:

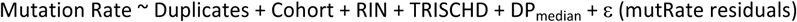

We understand *mutRate residuals* (*ε*) as the variability of the observed (raw) mutational rate, which is not explained by non-biological (technical) features such as PCR duplicate rates, cohort, or RIN. In order to assess the effect of age and tissue on mutation rates, we assessed the relation of the remaining variability (*mutRate residuals*) and the age of an individual at death, separately for each tissue, using a Spearman’s rank correlation test (all p-values were corrected with FDR).

### Mutational signatures

Mutational signatures were computed using the R package *deconstructSigs* (Rosenthal et al., 2016) and only signature weights greater than 0.1 were shown in plots.

For computing mutational signatures of embryonic mosaic mutations all individuals were considered for which at least 10 tissues were sequenced. For calculation of signatures of somatic mutations acquired during the lifespan only individuals older than 60 years were included in the analysis in order to increase the number of mutations related to mutagenic processes. Again, we focused on mutations found in exons due to the limited RNA-seq coverage in intergenic and introic regions. We obtained mutational signatures for each tissue separately, as well as for groups of tissues based on predominant environmental exposures, with a specific focus on:

- Sun-exposed skin
- Non-sun-exposed skin
- Exposure to mutagens in food: colon, esophagus-mucosa, small intestine, liver and stomach
- Brain tissues: Brain-Anterior_cingulate_cortex_BA24, Brain-Hippocampus, Brain-Substantia_nigra, Brain-Caudate_basal_ganglia, Brain-Cerebellar_Hemisphere, Brain-Frontal_Cortex_BA9, Brain-Spinal_cord_cervical_c-1, Brain-Amygdala, Brain-Cortex, Brain-Cerebellum, Brain-Hypothalamus, Brain-Nucleus_accumbens_basal_ganglia, Brain-Putamen_basal_ganglia

### Identifying signatures of positive selection in cancer genes using dN/dS

To estimate the extent of selection acting on somatic mutations in healthy tissues we used the SSB-dN/dS method (Zapata et al., 2018), which calculates the trinucleotide-corrected ratio of nonsynonymous to synonymous mutations from NGS data (Zapata et al., 2018). Somatic mutations identified by *RF-RNAmut* were annotated using Variant Effect Predictor (VEP). To increase statistical power, we only considered constitutively expressed genes having more than 5 TPM in at least 75% of patients for a focal tissue. We computed SSB-dN/dS in each tissue separately, and in the pan-tissues combinations listed above, using 192 parameters for nucleotide bias correction (correcting for mutation bias in all possible triplets on forward and reverse strand). However, we only computed dN/dS values for those tissues having at least 3 non-silent or silent somatic mutations in the analyzed genes. In addition to the exome-wide dN/dS provided in the output of the SSB-dN/dS method, we calculated the global dN/dS for 198 cancer genes (Martincorena et al., 2015) and 995 essential genes (Zapata et al., 2018). Finally, we focused on NOTCH1 and TP53 genes in order to replicate the findings of strong positive selection described recently (Martincorena et al., 2015, 2018; Yizhak et al., 2019; Yokoyama et al., 2019).

## Acknowledgement

We would like to thank the donors and their families for their generous gifts of organ donation for transplantation and tissue donations for the GTEx research study.

## Availability of data and materials

The data supporting the conclusions of this article were obtained from the Genotype-Tissue Expression (GTEx) portal and are available in the dbGaP repository, accession phs000424.v7.p2. Sequencing data of CLL individuals have been deposited at the European Genome-Phenome Archive (EGA, http://www.ebi.ac.uk/ega/), which is hosted by the EBI and the CRG), under accession number EGAS00000000092. Custom scripts used for somatic variant calling in this study are provided in https://github.com/Francesc-Muyas/RnaMosaicMutationFinder.

## Author Contributions

SO, RG and FM designed the study. FM developed bioinformatics methods and/or performed statistical analysis for mosaic mutation detection, mutational signature analysis, and estimation of mutation rates. LZ performed the evolutionary selection dN/dS analysis. FM and LZ generated the figures. SO and FM wrote the paper with the help of all authors.

