## Supplementary Information for "The rate and spectrum of mosaic mutations during embryogenesis revealed by RNA sequencing of 49 tissues"

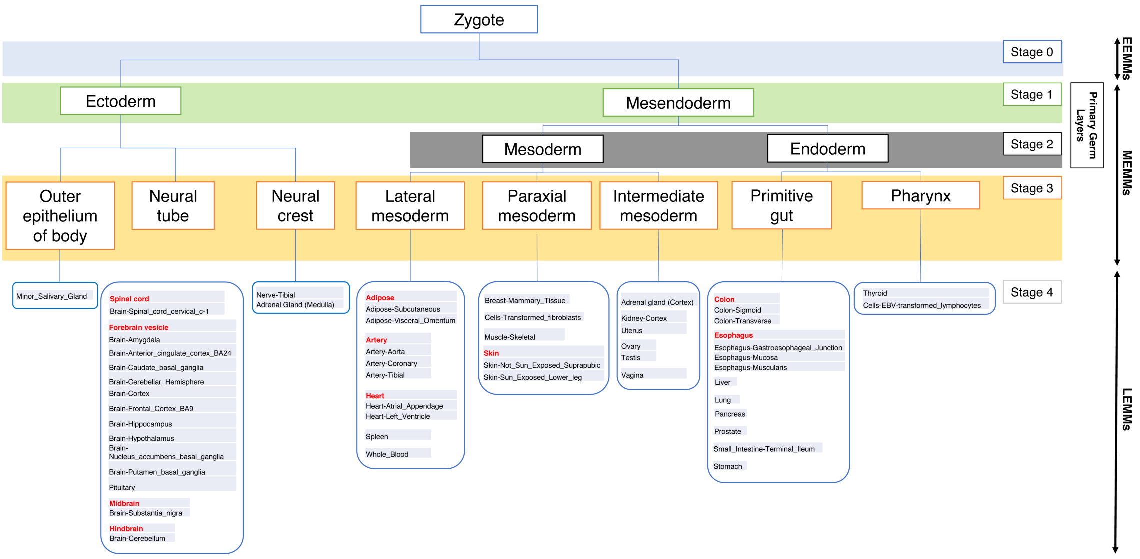


**Supp. Figure 1 | Lineage tree of human embryogenesis and organogenesis including 49 tissues studied in GTEx.** The *Stage* label groups the tissues based on critical phases of embryogenesis, approximately representing *Stage 0*: from first division of the zygote until late blastulation, *Stage 1*: late blastulation and implantation, *Stage 2*: gastrulation, *Stage 3:* neurulation, and *Stage 4:* organogenesis. Black arrows on the right represent the classification of embryonic mosaic mutations into early-embryonic mosaic mutations (EEMMs), mid-embryonic mosaic mutations (MEMMs), and late embryonic mosaic mutations (LEMMs), which has been used throughout this study.

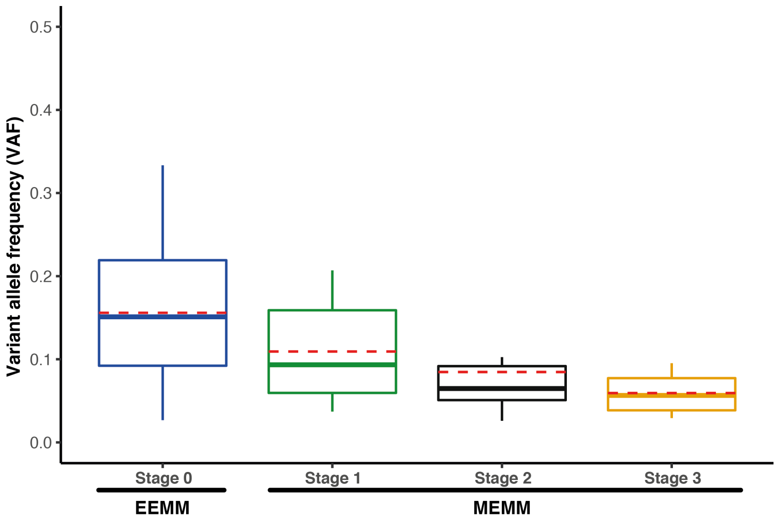


**Supp. Figure 2 | Variant allele frequency (VAF) distribution of mosaic variants mapped to different stages of embryogenesis** (see Supp. Figure 1). Early-embryonic mosaic mutations (EEMM) showed greater VAFs than mid-embryonic mosaic mutations (MEMM). The distinguishable stages of embryogenesis show a significant correlation with VAF when considering stages 0-3 described in Supp. Figure 1 (Spearman correlation (Rho) = -0.39 and p-value of $7.827 \times 10^{-5}$), as well as when considering only MEMMs, i.e. stages 1-3 (p-value = 0.0460 and rho = -0.3301).

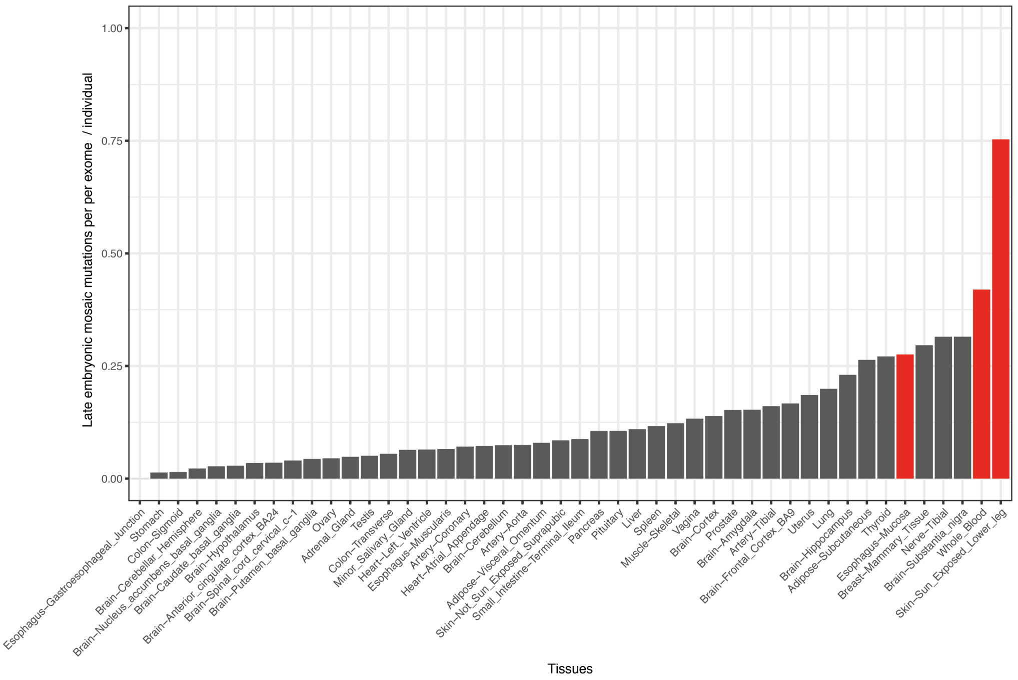


**Supp. Figure 3 | Rate of late embryonic (organ-specific) mosaic mutations observed per tissue and individual in human coding regions (45 Mbps).** Values were normalised by the number of informative samples per tissue. Red tissues were excluded in the identification of LEMMs, as they have been reported to harbour detectable clonal expansions.

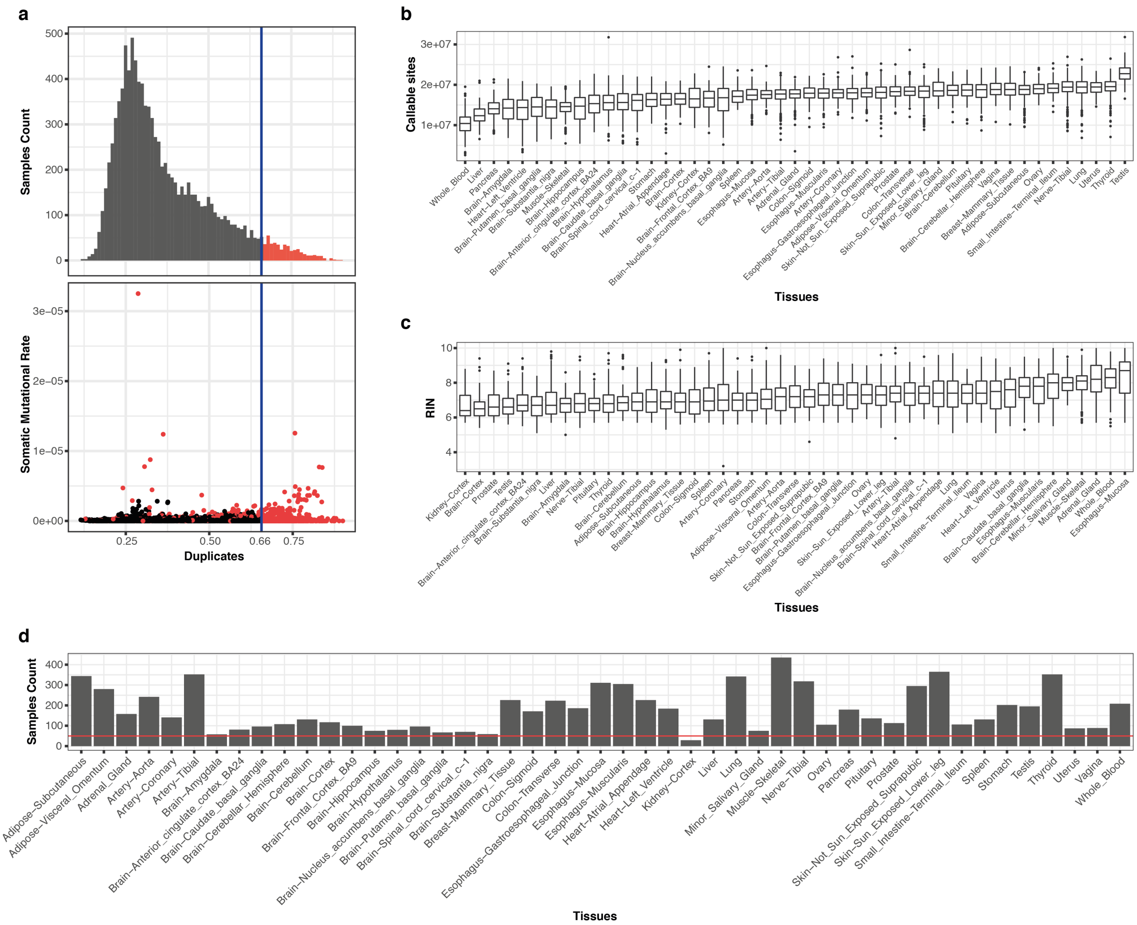


**Supp. Figure 4 | Quality control for RNA-seq data of the GTEx cohort for somatic mutation analysis.** (a) Samples with high PCR duplication rate have been excluded, as duplication rate positively correlates with the number of predicted somatic mutations (confounder), (b) number of callable sites per tissue is depending on the number of expressed genes per tissue, (c) Distribution of RIN values per tissue. (d) Number of sequenced individuals per tissue post QC filtering. Tissues with less than 50 samples (below red line) were removed from the study (only kidney).

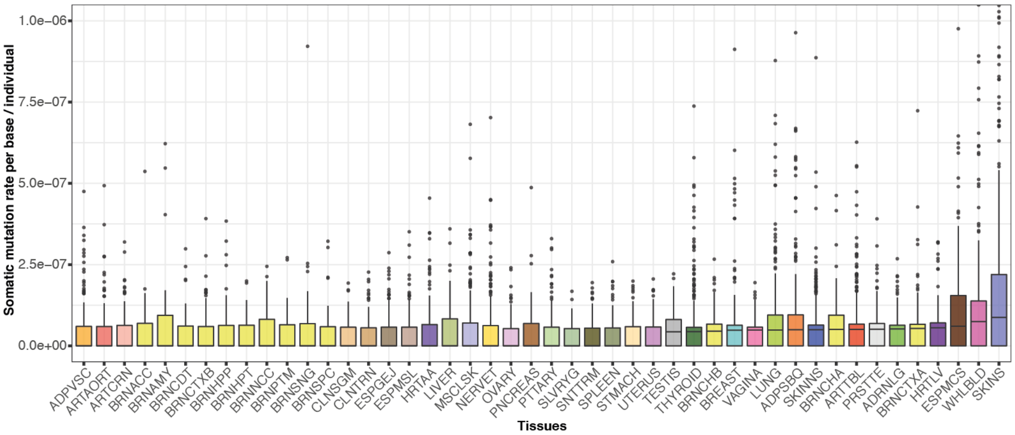


Supp. Figure 5 | **Uncorrected rate of somatic mutations per tissue, which we found to be confounded by technical biases such as PCR duplication rates, read coverage and RIN.** Residuals corrected for technical biases are shown in **Error! Reference source not found.**a.

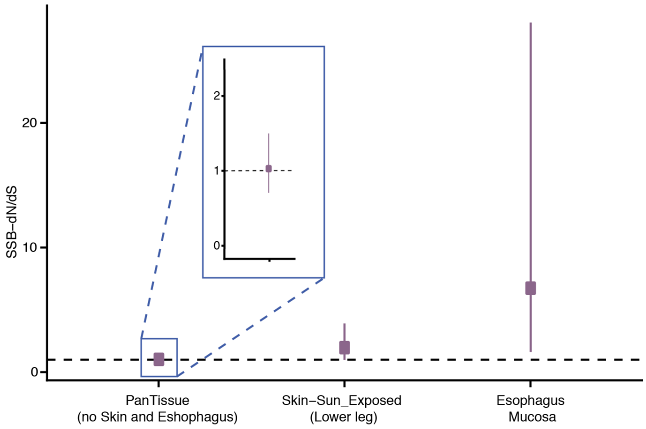


Supp. Figure 6 | **Signatures of positive selection in cancer genes identified for sun-exposed skin and esophagus-mucosa.** Pan-tissue analysis (without skin and esophagus-mucosa) revealed neutral evolution for the majority of tissues.

Supp. Table 1 | Performance of RNA-seq based variant detection in CLL samples using different thresholds for variant allele frequency (VAF).

| Mutation Type | VAF filter | Precision | Sensitivity | F1 |
|---|---|---|---|---|
| Somatic | 0.05 | 0.8511 | 0.5200 | 0.6456 |
| Somatic | 0.10 | 0.8511 | 0.6290 | 0.7234 |
| Somatic | 0.15 | 0.8511 | 0.7091 | 0.7736 |
| Somatic | 0.20 | 0.8511 | 0.7400 | 0.7917 |
| Somatic | 0.25 | 0.8511 | 0.7347 | 0.7886 |
| Germline | 0.25 | 0.9534 | 0.8600 | 0.9043 |

Supp. Table 2 | Number and rate of EEMMs and MEMMs in the four sets of constitutively expressed genes.

| Gene Selection | Num. Genes | Region Size | Num. Individuals | Num. EEMMs | EEMM Rate per Base and Individual | EEMMs per Exome and Individual* | Num. MEMMs | MEMM Rate per Base and Individual | MEMMs per Exome and Individual* |
|---|---|---|---|---|---|---|---|---|---|
| TPM >= 5 | 7,630 | 12,300,858 | 526 | 58 | 8.96E-09 | 0.4034 | 36 | 5.56E-09 | 0.2504 |
| TPM >= 10 | 5,231 | 7,660,529 | 526 | 31 | 7.69E-09 | 0.3462 | 19 | 4.72E-09 | 0.2122 |
| COV >= 20 | 6,888 | 11,078,846 | 526 | 48 | 8.24E-09 | 0.3707 | 30 | 5.15E-09 | 0.2317 |
| COV >= 30 | 5,370 | 8,300,031 | 526 | 33 | 7.56E-09 | 0.3401 | 22 | 5.04E-09 | 0.2268 |

Supp. Table 3 | Signature of selection in cancer genes. dN/dS values above 1 indicate positive selection.

| **Tissue** | **Ensembl ID** | **Hugo symbol** | **Total Number Mutations** | **Non-silent variants** | **Synonymous variants** | **dN/dS SSB** | **pval SSB** |
|---|---|---|---|---|---|---|---|
| Esophagus-Mucosa | ENST00000277541 | *NOTCH1* | 17 | 17 | 0 | 8.4617 | 0.0063 |
| | ENST00000269305 | *TP53* | 10 | 10 | 0 | 4.5655 | 0.0590 |
| Skin-Sun_Exposed_Lower_leg | ENST00000269305 | *TP53* | 11 | 10 | 1 | 2.8516 | 0.1035 |
| | ENST00000277541 | *NOTCH1* | 5 | 5 | 0 | 4.0133 | 0.0826 |
| Pan Tissue (except Skin and Esophagus) | Cancer Genes | | 140 | 103 | 37 | 1.0295 | 0.8800 |
